# Accurate informatic modeling of tooth enamel pellicle interactions by training substitution matrices with Mat4Pep

**DOI:** 10.1101/295857

**Authors:** Jeremy A. Horst, Jong Seto, Ersin Emre Oren, Orapin V. Horst, Ling-Hong Hung, Ram Samudrala

## Abstract

**Motivation:** Protein-hydroxyapatite interactions govern the development and homeostasis of teeth and bone. Characterization would enable design of peptides to regenerate mineralized tissues and control attachments such as ligaments and dental plaque. Progress has been limited because no available methods produce robust data for assessing phase interfaces.

**Results:** We show that tooth enamel pellicle peptides contain subtle sequence similarities that encode hydroxyapatite binding mechanisms, by segregating pellicle peptides from control sequences using our previously developed substitution matrix-based peptide comparison protocol (Oren et al., 2007), with improvements. Sampling diverse matrices, adding biological control sequences, and optimizing matrix refinement algorithms improves discrimination from 0.81 to 0.99 AUC in leave-one-out experiments. Other contemporary methods fail on this problem. We find hydroxyapatite interaction sequence patterns by applying the resulting selected refined matrix (“pellitrix”) to cluster the peptides and build subgroup alignments. We identify putative hydroxyapatite maturation domains by application to enamel biomineralization proteins and prioritize putative novel pellicle peptides identified by In stageTip (iST) mass spectrometry. The sequence comparison protocol outperforms other contemporary options for this small and heterogeneous group, and is generalized for application to any group of peptides.

**Availability:** Software to apply this protocol is freely available at github.com/JeremyHorst/Mat4Pep and compbio.org/protinfo/ Mat4Pep.

**Supplementary information:** Available at *Bioinformatics* online.

## 1 Introduction

Mechanisms of protein to hydroxyapatite interactions in tooth and bone remain elusive. Here we introduce a generalized approach for detecting patterns in peptide sequences, and apply the method to describe amino acid sequence features that may control interactions with forming and mature hydroxyapatite.

The enamel pellicle is a layer of peptides derived from saliva that binds directly to and coats tooth enamel, and is bound by early colonizer dental plaque bacteria. Sequences for the enamel-binding peptide constituent of the human enamel pellicle (pellicle peptides) have been described (Siqueira et al., 2007; Vitorino et al., 2007; Vitorino et al., 2008; Siqueira and Oppenheim, 2009). The greater context of the salivary proteome, from which these proteins arise, has been explored and tapped as readily available diagnostic samples, for example to detect cancer (Hu et al., 2008).

Physiologic details of enamel binding have been explored to the extent of measuring adhesion strength for the saliva-derived enamel pellicle to oral bacteria (Mei et al., 2009). Specific peptides have been designed to replace this pellicle handle by which oral flora adhere to the tooth (Li et al., 2009). Yet the mechanisms of peptide to enamel adhesion are still poorly understood.

One type of hydroxyapatite interaction is obvious from clues in nature. Comparison of the aspartate-serine-serine (DSS) repeats in dentin phosphoprotein (DPP) to the hydroxyapatite unit cell hints at a template of carboxylates interacting with calcium and hydroxyls interacting with phosphates. Similar or enhanced affinities are observed upon mutation to residues bearing the same functional groups but different side chain lengths (Yarbrough et al., 2010).

Relatively few proteins directly interact with tooth and bone hydroxyapatite. Besides a couple proteins like DPP, domains responsible for direct hydroxyapatite interactions are sparsely characterized. No atomic resolution structures of proteins that physiologically interact with hydroxyapatite are available, except osteocalcin (PDB entry 1q8h), so structural analysis for these proteins is elusive. Neither the DSS repeats of DPP nor the γ-carboxy glutamic acids of osteocalcin are present in the pellicle peptides or enamel-forming proteins, so no homology-based inferences are accessible.

While no obvious similarities are found among the pellicle peptides (Siqueira and Oppenheim, 2009), this set of 78 peptides from 29 proteins comprises the largest and most diverse information on hydroxyapatite interactions. We hypothesize that patterns in the sequences of enamel pellicle peptides can drive discovery of protein-hydroxyapatite interactions and mechanisms.

We anticipate that the mechanisms underlying peptide-hydroxyapatite interactions produce nontrivial similarities in the protein sequences that can drive the training of a sequence comparison algorithm to successfully discriminate enamel-binding pellicle peptides from control sequences. However, physiologic peptides that do not bind tooth enamel have not been directly observed, so we fabricate decoy sets as the negative control instances to feed the algorithm. The regions of the source protein sequences least likely to bind enamel are those areas from which the pellicle peptides are not derived - they are exposed to the same environment that enables enamel interactions and therefore it is likely that they would be observed if they did bind enamel. We derive the decoy control set from these protein regions. Omission by lack of observation is not sufficient evidence to identify absent function (enamel binding), but discrimination from pellicle peptides would give evidence for differential evolution and validate the approach.

Previously we exploited the sequence similarities of phage display peptides that bind to inorganic surfaces to program an amino acid substitution matrix, and subsequently designed peptides with enhanced binding affinity to that surface (Oren et al., 2007).

Although the pellicle set has amino acid content patterns (Figure 1), there are not sufficient position-specific patterns to enable construction of a multiple sequence alignment as necessary for application of commonly used sequence comparison algorithms such as PSI-BLAST or hidden-Markov models (HMMs). Nor are neural networks able to perform better than random in leave-one-out experiments (SciKit-Learn; Figure S1). The Needleman-Wunsch algorithm does not require a strong pairwise alignment to construct a comparison, and thus may capture more diffuse sequence similarities, as in a heterogeneous set of enamel binding peptides.

**Fig. 1.**
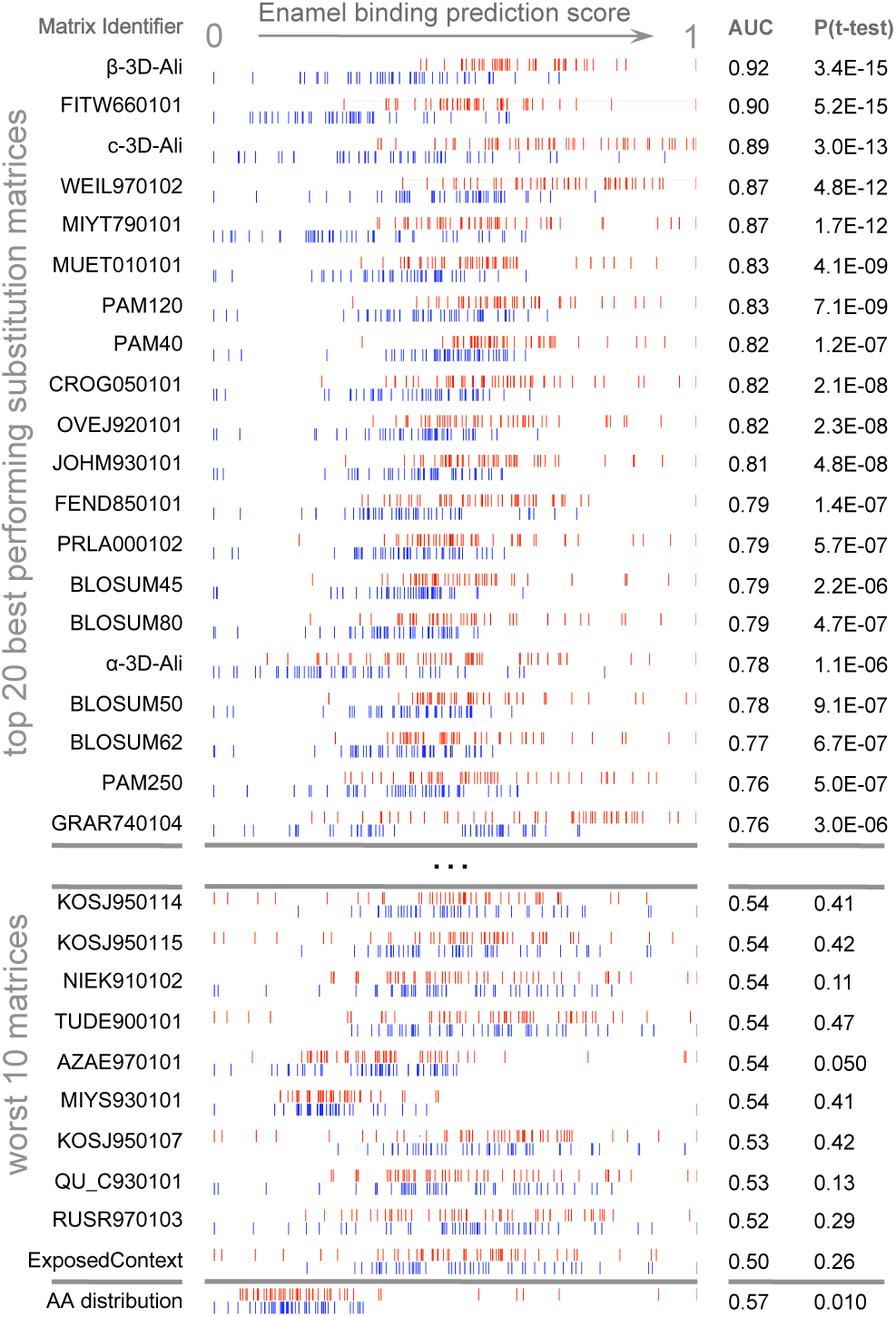
Discrimination of enamel pellicle peptides. The scoring of 49 pellicle peptides (red) from 49 control sequences (blue) in a modified leave-one-out experiment is shown for amino acid content, and the top 20 and worst 10 performing substitution matrices. Each row represents application of one matrix, for which normalized scores are plotted for each pellicle and control sequence. Better discrimination is seen at top, with pellicle peptides assigned higher scores (red to the right) and controls assigned lower scores (blue to the left). No overlap for the profiles of pellicle and control markers would indicate perfect discrimination. Most matrices discriminate more accurately than amino acid content (at bottom), demonstrating the importance of the sequential and spatial arrangement of residues.

The Needleman-Wunsch dynamic programming algorithm finds the optimal global alignment for two protein sequences with respect to the scoring system being used (Needleman and Wunsch, 1970), which includes a substitution matrix and penalties for opening or extending gaps in the alignment. The more popular Smith-Waterman algorithm is essentially a variant of the Needleman-Wunsch algorithm with zeroed negative matrix values, such that local alignments are optimized (Smith and Waterman, 1981).

Optimal gap penalties are found using a simple grid search. Finding the optimal matrix values by which to score the potential alignment of two sequences is the challenge (Kawashima et al., 2008). The combination of 39 integer values (from −19 to 19) for each of the 210 possible amino acid substitutions in a symmetric matrix, 39^210^, is too many to enumerate (39^400^ if asymmetric). Substitution matrices can be calculated directly by comparative analysis between sets, but alignments must already be known. Unless the set is large enough to represent the relevant evolutionary relationships, this approach has the propensity to become too specific to the data set, i.e. overtraining.

One technique that performed well for the phage display-derived inorganic surface-binding problem was to exploit a substitution matrix calculated with a widely diverse set of proteins (e.g. BLOSUM62, PAM250) and refine the values to the dataset (Oren et al., 2007). Refinement may not resolve to a near optimal matrix, as coarse integer-based scoring functions result in local maxima and weak trajectories to guide improvement. Therefore, here we sample many starting matrices from the diverse set in AAindex (Kawashima et al., 2008). In this work we ask whether a sequence analytic algorithm can select and refine a substitution matrix to discriminate functional peptides of dissimilar lengths from controls, find these peptides from within their source proteins, and identify mechanistic patterns in these natural sequences.

## 2 Methods

### 2.1 Data sets

#### Acquired enamel pellicle peptides

The peptides taken to be true pellicle constituents in this work are the 78 from 29 salivary proteins observation in multiple studies and described by Siqueira and Oppenheim in 2009. For use in our bioinformatic experiments, we aligned the peptide sequences, removed 100% redundant sequences, and combined overlapping portions from the same protein. The resulting new pellicle peptide fragment set includes 49 peptides, 8-36 residues in length (Supplemental Table 1).

#### Control sequences

For controls in training and back-testing we used fragments of the 29 proteins not observed within the 78 acquired enamel pellicle peptides. We retrieved random fragments matching the number and length of the peptides, in regions not overlapping the pellicle peptide sequences. When intervening stretches were not abundant or long enough to derive a matching set, we retrieved additional fragments from random other proteins in the set. The resulting decoy control set includes 49 peptides, 8-36 residues in length (Supplemental Table 2).

#### Additional negative sequences from other proteins

To increase information content for matrix training and enhance relevance to nonpellicle proteins, we derived additional presumed nonfunctional sets matching the pellicle peptide set in length and quantity. One set was produced by extracting random parts of any human protein secreted in the saliva (Supplemental Table 3). Additional sets were constructed from random sequences by combination of amino acids selected to mimic the composition in UniProt (The UniProt Consortium, 2007; Supplemental Table 4). We attempted training with and without each of the additional background sequence sets. Additional negative sequences were included as controls during training and not assessment. Wherever use of these sequences did not disrupt training, they were included to enhance relevance to other proteins.

### 2.2 Training protocol

#### Similarity calculations

The total similarity score function (TSSF) is the primary output metric used to differentiate between pellicle peptides and control sequences. Matrices, gap values, and training paths were optimized by maximizing TSSF. The TSS is applied as the sum of Needleman-Wunsch scores (Needleman and Wunsch, 1970) for all alignments between two sets, normalized by peptide length and the number of sequences in each set (Oren et al., 2007). Previously we used the difference of TSS for functional peptides to themselves (TSS.ff) and functional to non-functional peptides (TSS.fn; TSSF = TSS.ff – TSS.fn; Oren et al., 2007). Here we considered TSS for non-functional to themselves (TSS.nn) and non-functional to functional TSS (TSS.nf), as the difference (TSSF = TSS.ff + TSS.nn-TSS.fn-TSS.nf) or the quotient (TSSF = TSS.ff * TSS.nn / (TSS.fn * TSS.nf)). We also attempted training to maximize the difference between the third lowest (to allow for outliers) scoring pellicle peptide and the third highest scoring control sequence.

#### Gap penalties

Gap penalties were trained by selecting the maximal score in an integer grid based search [−16, −1] for the gap open penalty and [−8, −1] for the gap extend penalty. Gap penalties were only trained before altering substitution matrices, and not iteratively, due to their potential volatility during a training process.

#### Amino acid substitution matrices

We took starting matrices from 75 amino acid substitution matrices in AAindex (Kawashima et al., 2008). Matrix elements are perturbed as integers within the range −19 to 19.

#### Refinement paths

We evaluated three substitution matrix refinement paths. We perturb the starting matrix values by either greedy or modified Monte Carlo trajectories. The greedy algorithm considers all possibilities and then chooses the path that makes the largest magnitude of improvement (increased TSSF). We also attempted either local maximization by using the minimum unit of the matrix, or a modified Monte Carlo search for the global maximum by using a random value less than the maximum difference in the matrix, with the decision of keeping each sequential step made after local maximization. We also attempted refinement paths wherein the importance of query versus data set amino acid and overall trends in amino acid type were simultaneously examined, rather than amino acid type combinations (e.g. the target position being an alanine versus both query and target being alanine), as all sequential combinations of mutating columns, rows, and cells of the matrix. Refinement paths were followed until changes no longer resulted in improvements. Monte Carlo refinement was stopped after five consecutive attempts failed to make an improvement.

### 2.3 Assessment

#### Leave one protein out experiments

We attempted to discriminate pellicle peptides from control sequences by total similarity score (Figure 1). To assess accuracy, we performed modified leave-one-out experiments: while scoring a peptide we remove all sequences (pellicle peptides and controls) from the same protein. A normal leave one out experiment involves removing one constituent from the set, training on the rest, scoring the constituent, and repeating for each instance. Here peptides are separated by protein such that in the benchmark the algorithm never learns from and applies information to peptides from the same protein, because sequences in the same protein are likely to contain mutual information.

#### Statistical metrics

The receiver operating characteristic (ROC) compares sensitivity (true positives) across all ranges of specificity (true negatives; Figure 2a). The precision recall curve compares the precision at all ranges of recalled selections (Figure 2b). The Matthews correlation coefficient (MCC; Matthews, 1975) measures the correlation of true positives, false positives, false negatives, and true negatives. The MCC curve plots this correlation across a range of thresholds (e.g. 0.01 steps from 0 to 1) for indicating a true or positive result (Horst, 2010). The complexity of a MCC curve informs the capacity for improvement by further training, and identifies the threshold cutoff score that results in the most informative predictions (Figure 2c). Area under the ROC curve (AUC) and one-tailed unpaired unequal variance Student’s T-test (p values) tested significance.

**Fig. 2.**
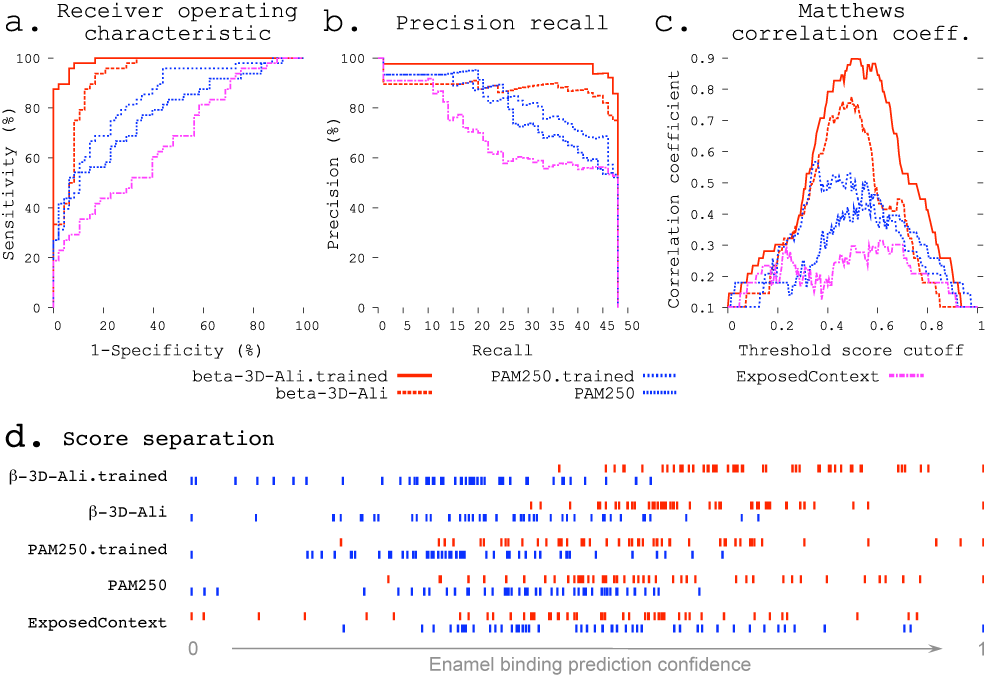
Refinement improves enamel pellicle peptide discrimination in a modified leave-one-out experiment. The β-3D-Ali.trained (solid red line, see key below panels a-c) and PAM250.trained (blue coarsely dashed line) matrices demonstrate increased predictive ability across three rigorous metrics from the β-3D-Ali (red dashed line) and PAM250 (blue thinly dashed line) matrices, respectively. Comparison is given to the worst performing matrix (ExposedContext = KOSJ950113). a. Receiver operating characteristic curve. b. Precision recall curve. c. Matthews correlation coefficient (MCC) curve. The complexity of each MCC curve informs the capacity for improvement: the untrained matrices both show a large local minimum, lost with improvement in the correlate trained curve. d. Score distributions (as in Figure 1) show greater separation of pellicle peptide (red) and control sequence (blue) scores after training.

#### Amino acid content calculation

To evaluate whether sequential orientation (position) influences enamel binding, we assessed whether the accuracy of scoring each amino acid in a query peptide by the proportion of the amino acid type in pellicle peptides versus controls.

### 2.4 Application to full protein sequences

We evaluate the ability to recapture pellicle regions from full protein sequences by generating a score for each residue in the protein, considering the surrounding region. We applied the sliding window approach for each unique length of pellicle peptides. For this problem, it is uncertain whether it would be better to choose segments of one particular length, or to exhaustively create segments of all pellicle peptide lengths. Even then, it is not known how to consider the similarity scores for the various segments to which each particular residue contributes. For both a single window length (the median of all peptide lengths) and enumeration of the lengths, we evaluated the application of the mean of the similarity scores for overlying segments and the maximum score for each. Maintaining consistent fragment lengths between query and comparison sets avoids a difficult normalization problem. We compared the predictive ability of residue scores to recapture the pellicle peptides from the entire protein sequences, again using the leave one protein out approach (Figure 3).

**Fig. 3.**
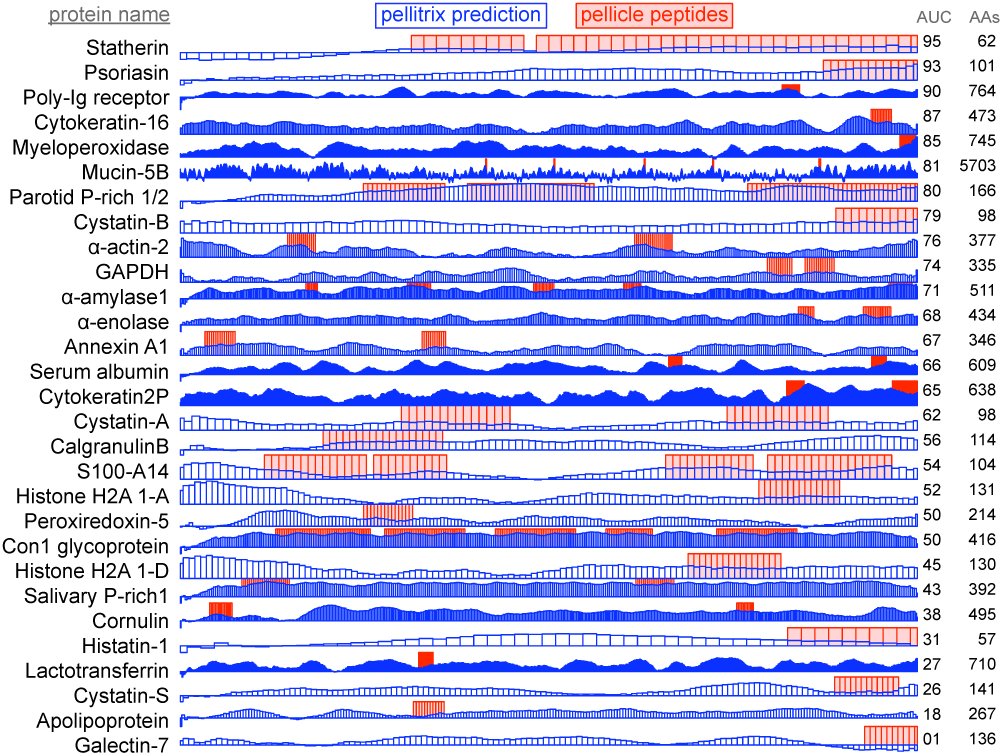
Enamel pellicle peptide recapture from complete proteins. Predictions of enamel affinity by the refined β-3D-Ali matrix (pellitrix) for each residue are plotted in blue for each enamel pellicle protein. Scores represent the mean of the similarity scores between all peptides derived from other proteins (modified leave one out experiment) and all possible overlapping sequence fragments of lengths matching the pellicle peptides (sliding window fragmentation). Experimentally derived pellicle peptides are shown as red blocks. Overlap of high blue bars with the red blocks denotes recapture of pellicle peptides from the parent protein. Protein length (AAs) and per-residue recapture accuracy (AUC) are listed at right.

### 2.5 Cluster analysis

To study the sequence patterns identified in training, we derived sequence clusters by analyzing the network of comparisons between all enamel pellicle peptides using the best selected and refined matrix. We filtered the resulting similarity scores by the threshold cutoff that gave maximum information in the benchmark according to the MCC plot (Figure 2c). We then input the supra-threshold similarities as force vectors into a clustering algorithm. We depicted the resulting network using cluster analysis in Cytoscape (Shannon et al., 2003). Subcluster networks were identified from the graph, and aligned by CLUSTALW (Larkin et al., 2007) using the same substitution matrix (Figure 4).

**Fig. 4.**
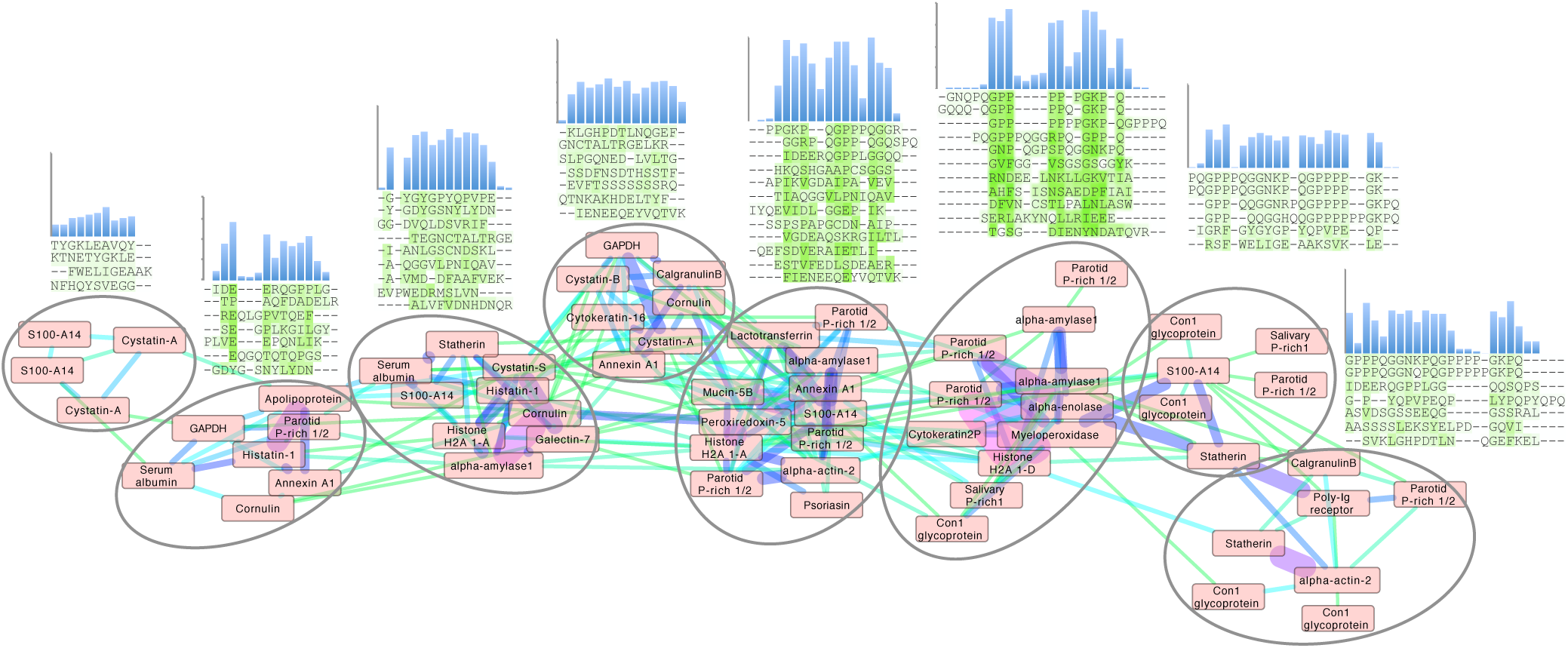
Cluster analysis of the enamel pellicle peptide sequence similarity network. Shown is the relation of the 78 peptide sequences (nodes) clustered with edge weights given by the trained β-3D-Ali matrix (pellitrix) similarity scores. The initial network was generated from an all-against-all matrix of these scores, for which edges were defined as any similarity score above the threshold cutoff corresponding to the maximum Matthews correlation coefficient in Figure 2c. The magnitude of similarity between pairs of peptides is shown as increasing from green to violet (edges). Protein names that appear multiple times indicate alternate peptides derived from the same protein. Node placement was adjusted slightly to enable viewing of protein names. The multiple sequence alignments display trends for each subcluster (circles), which suggest residue patterns for stabilizing extended beta strand and polyproline helix conformations; mediating calcium interactions by adjacent carboxyl and amide residues; mediating phosphate interactions by alternating hydroxyl moieties; and pi interactions and/or hydrophobic exclusion by aromatic moieties.

### 2.6 Software

All code was written in Python version 2.7. The Needleman-Wunsch algorithm implemented as ggsearch35 was taken from the FASTA suite version 35.4.11 (Pearson and Lipman, 1988). Statistical tools employed in the assessment were written locally and extensively checked against both SPSS and STATA. Figures were depicted with GNUplot (Williams et al., 2012; gnuplot.info) and R (R Core Team, 2017).

### 2.7 Pellicle peptide characterization

#### Sample collection

De-identified samples were collected with consent under UCSF IRB exempt protocol, as previously (Siqueira and Oppenheim, 2009). Briefly, two hours after prophylaxis with pumice, and limitation from eating, teeth were rinsed with sterile deionized water, and scraped with micropipette tips, which were vortexed in 10 mM PBS, and pooled.

#### In stageTip (iST) mass spectrometry

Samples were moved into urea lysis buffer, treated with trypsin/lysC, reduced with TCEP, and alkylated with 2-chloroaceatmaide in a “single pot” system to minimize sample loss and contamination, then placed in 0.1% acetic acid and 80% acetonitrile until LC/MS mass spectrometry (Thermo Scientific LTQ-Orbitrap Velo, ThermoFisher Scientific), as previously (Kulak et al., 2014).

#### Peptide data analysis

MaxQuant and Perseus (Cox and Mann, 2008) were applied to identify and assess the validity of source protein sequences for each observed peptide amidst the human proteome.

## 3 Results

### 3.1 Selected and refined peptide discrimination

We demonstrate the ability of the matrix sampling and refinement protocol to optimize performance in discriminating pellicle from control sequences (Figures 1 and 2). Three statistical metrics verify marked improvement of two highly different substitution matrices (Figure 2). The β-3D-Ali matrix (MEHP950102) was selected for optimal peptide discrimination, and refined from 0.92 (p=3.4*10^-15^) to 0.99 AUC (p=3.4*10^-26^). We present the optimized substitution matrix and values changed during training in Supplemental Table 5. The PAM250 matrix (DAYM780301) was refined from 0.76 (p=5.0*10^-7^) to 0.84 AUC (p=4.5*10^-10^). We extended the refined β-3D-Ali matrix (“pellitrix”) to estimate the likelihood of any single residue binding tooth enamel and calculated recovery of the pellicle peptides (0.75 AUC; Figure 3). We analyzed pellicle peptide similarities with the refined selected matrix to gain mechanistic insight into pellicle-enamel interactions (Figure 4). Finally, we applied pellitrix to predict biomineralization interactions in enamel matrix proteins (Figure 5), and to prioritize novel peptides observed in the enamel pellicle (Figure 6).

**Fig. 5.**
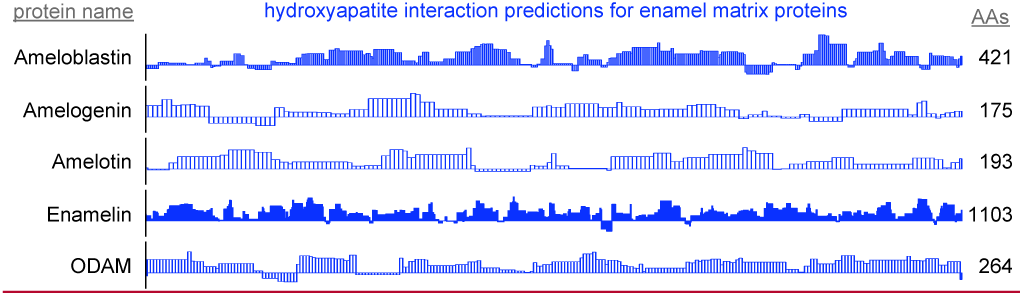
By-residue likelihood of hydroxyapatite interactions for enamel matrix proteins. The refined selected matrix was applied to find the similarity of the region surrounding each residue to the enamel pellicle peptides. Scores are normalized to the highest and lowest scores observed for all peptides and control sequences. Length of proteins is shown at right. High scoring regions likely correspond to functional areas that interact with mature or maturing enamel. Low scoring areas may carry out functions not consistent with mature enamel, such as hydroxyapatite nucleation and endoprotease cleavage.

**Fig. 6.**
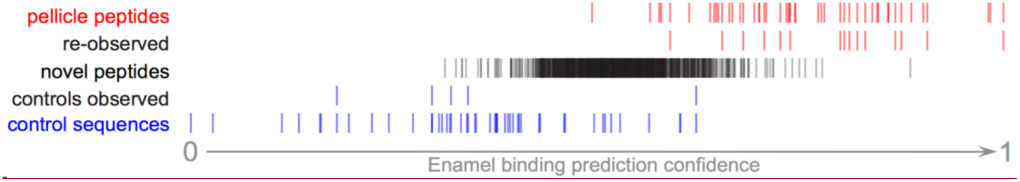
Pellitrix scores for novel peptides observed in the enamel pellicle by iTS mass spectrometry. 15 pellicle peptides (re-observed) and 5 control sequences (controls observed) occurred within 1,265 sequences. Scores for the remaining sequences (novel peptides) are plotted in context. The 92 peptides with scores above the range of control sequences are likely to contribute physiologically to the enamel pellicle (Supplemental Table 9).

### 3.2 Matrix sampling

AAindex (Kawashima et al., 2008) matrices discriminated pellicle peptides from control sequences with performance ranging from discriminating the majority of pellicle peptides, to none (Supplemental Table 6). Figure 1 shows the distribution of scores for pellicle peptides and control sequences for the top twenty matrices, the worst ten, and scoring by amino acid content. The β-3D-Ali matrix most accurately separated pellicle peptides from controls, and along with the PAM250 matrix was used for further analysis.

### 3.3 Matrix refinement

The refinement protocols improved performance on the task of sorting pellicle peptides from control sequences for both the PAM250 and β-3D-Ali matrices (Figure 2, Supplemental Table 7).

#### Similarity calculations

All three subtraction-based similarity calculations resulted in improvement for the PAM250 and β-3D-Ali matrices, whereas the quotient based similarity calculation did not. The most significant improvements in the matrices arose consistently from including the relation of control sequences to themselves and to the pellicle peptides in the total similarity score (TSSF = TSS.ff + TSS.nn - TSS.fn - TSS.nf).

#### Refinement paths

The best and most consistent matrix refinement protocol was achieved by a greedy path, exhausting improvements from changing all values in each column together, exhausting improvements similarly in the rows, then optimizing whole columns and rows with the modified Monte Carlo search. The greedy algorithm uses more processor time than a random or Monte Carlo path, as both positive and negative trajectories for each position must be considered before progressing to the next step. Each training combination reaches completion in 4 hours on a 4.8 GHz processor (~10,000 pairwise comparisons per minute).

The order of starting permutations with matrix row (query amino acid type) or column (pellicle / control amino acid type) affected the performance of the matrix. Only a few random paths starting with rows improved performance, while many training conditions improved accuracy when starting with columns. Adding Monte Carlo perturbations of columns and then rows as a last set of steps after the described greedy path improved performance in nearly all cases, whereas Monte Carlo perturbations of the cells never did.

#### Training data set combinations

Inclusion of the additional background sequences into the controls improved the discriminatory performance of both PAM250 and β-3D-Ali matrices slightly (AUC ~1%) with statistical significance (p<0.01).

#### Relationship of improvement to matrix distance

Across all matrices, the magnitude of improvement ranged from 0.002 to 0.41 AUC, with many nearing perfect discrimination. The arithmetic distance between the matrices before and after training correlated to improvement (Pearson’s R=0.55; Figures S2-S4).

#### Preferences of the trained matrix

Pairwise amino acid substitution scores for the identical residue and for the mean of all possible residue substitutes indicate the importance of matching each particular amino acid type in the final selected and trained matrix (Supplemental Table 8). For example, it is preferred that glutamic acid is aligned with another glutamic acid (score = 2.00), but self match is penalized for leucine (−1.40) and arginine (−2.00).

### 3.4 Protein binding region recapture

Accuracy of pellicle peptide recapture from full protein sequence depended largely on the formalism. Comparing protein segments of the median pellicle peptide length (14 residues) with pellitrix achieved 0.75 AUC for the mean score, and 0.54 AUC for the maximum. A similar difference was found for enumerating all lengths: 0.69 AUC for the mean and 0.54 AUC for the maximum. A caveat to this experiment should be noted: while the leave-one-out design avoids comparing peptides directly to any part of their source protein sequence, the information trained into the matrix in the selection and refinement steps cannot be removed and so biases this experiment. Without training, the β-3D-Ali matrix achieves 0.73 AUC using the mean of the multiple sliding windows, again the highest of all matrices (Supplemental Table 7).

### 3.5 Pellicle peptide sequence cluster analysis

Application of pellitrix to compare all 78 pellicle peptides to each other resulted in a network of context-specific sequence similarities (Figure 4). Multiple sequence alignments constructed with pellitrix illustrate in each column the amino acid types that can function similarly within the specific context of protein-hydroxyapatite interactions.

### 3.6 Novel pellicle peptide prioritization

1,265 unique peptides from the pooled pellicle sample were observed at least twice by iST secondary mass spec (MS/MS), identified by MaxQuant, and judged as significant by Proteus (Supplemental Table 9). Figure 6 shows that the range of pellitrix scores for these peptides falls within that of pellicle peptides and control sequences. The mean score falls at the center of the range (0.52) and the highest control sequence score corresponds to 1.5 standard deviations from the mean for the novel peptides. 15 of the 49 pellicle peptides and 5 control sequences were observed.

## 4 Discussion

### 4.1 Advancement in biomineralization

The ability of many amino acid substitution matrices to accurately discriminate enamel pellicle peptides from control sequences (Figure 1) demonstrates the presence of discernable sequence patterns, which likely underlie the common function of enamel hydroxyapatite binding. Cluster analysis (Figure 4) suggests peptide groups likely to share similar mechanisms and sequence patterns to facilitate them. The refined selected matrix can be used to analyze sequences for likelihood of contributing protein-hydroxyapatite interactions in peptides (Figure 2,6), whole protein sequences (Figure 3,5), and to design novel peptides.

Novel peptides may be designed with controllable binding affinities, used as a supplementary pellicle coat to control the attachment of oral flora, or as an adjuvant vehicle for controllable delivery of saliva replacements such as anticariogenic antibiotics or remineralizing agents (Yarbrough et al., 2010).

### 4.2 Advancement in bioinformatics

The improvements we introduce to our protocol to develop peptide similarity detection tools increased the final trained matrix discriminatory ability from 0.81 AUC with the old protocol to 0.99 AUC with the new protocol. Meanwhile, standard sequence comparison methods failed on this problem (Figure S1). MCC plot analysis indicates the training of this matrix has approached saturation (Figure 2c). The most significant improvements arose from sampling many starting substitution matrices, incorporating all peptide and control comparisons into the total similarity scores, and Monte Carlo optimization of columns and rows after greedy refinement. This approach may be able to learning patterns in any group of functional peptides, and is available as software called Mat4Pep for use and development.

### 4.3 Matrix sampling

Discriminatory performance across the matrices may indicate relevance to the context for which the matrix was calculated. Matrices built for general protein sequence comparison exhibited intermediate performance. The best performance came from a matrix built specifically to align β-strands in 38 3D-Ali protein structure families (Mehta et al., 1995), while matrices derived in parallel from random coils performed third, and that for α-helices ranked 16^th^. These secondary structures match observations that regions which interact with hydroxyapatite adopt beta strand or polyproline type II extended conformations (Jin et al., 2009; Carneiro et al., 2016).

### 4.4 Protein binding region recapture

Application of scores to the derivative proteins (Figure 3) shows successful modeling of a significant subset of enamel binding mechanisms. High scoring regions at locations where pellicle peptides have not been measured are predictions of areas that may bind enamel, for example the amino terminal regions of α-actin 2, cystatin-A, S100-A14, histone H2As 1-A and 1-D (Figure 3).

Recapture of pellicle peptides from whole protein sequences is better than average for 21 of 29 proteins, with a by-residue AUC of 0.75 across all proteins. Poor performance of the PAM250 matrix (AUC=0.31) highlights the uniqueness of sequence traits within these peptides of such rare function, and therefore the importance of using similarity matrices with maximal relevance to any particular group of proteins under study. This analysis demonstrates novel ability to understand, predict, and potentially design protein to hydroxyapatite interactions.

### 4.5 Pellicle peptide sequence cluster analysis

Each cluster in the network analysis displays trends in multiple sequence alignments (Figure 4). We observe tolerance for swapping residue identity but maintenance of chemical moieties: adjacent carboxyl or amide residues may facilitate calcium interactions (Horst and Samudrala, 2010), and alternating hydroxyl moieties may mediate phosphate interactions. Stretches of prolines may stabilize extended conformations, facilitating surface interactions. Proline almost never aligns with glutamine, suggesting noninterchangeable roles for the two most abundant residues in these peptides. Residue types most commonly involved in enzymatic catalysis (in order: EKDHRSTCYNQAFGMLWIVP; Wang et al., 2008) are seldom aligned with identical amino acid types in these clusters. These patterns suggest greater structural conservation with variance allowed for chemical interactions, which fits the presentation of calcium and phosphate on hydroxyapatite.

### 4.6 Application to enamel matrix proteins

High scoring regions in five enamel matrix biomineralization proteins (Figure 5) are predicted to participate physiologically in enamel development. Low scoring areas may carry out functions that require staying away from mature enamel, such as mineral nucleation or cleavage by endoproteases (Horst, 2010). These data may be used to derive peptides, or inform mutation experiments to drive mechanistic understanding of enamel development.

Predictions of hydroxyapatite interactions in Amelogenin (Figure 5) coincide with experimental hydroxyapatite binding data for peptides derived from the Amelogenin sequence (Gungormus et al. 2012). This convergence emphasizes the validity of the protocol in finding the enamel-binding regions in related proteins.

### 4.7 Novel pellicle peptide prioritization

Recent advances in mass spectrometry protocols and technology motivated re-assessment of pellicle peptides. Observation of 15 known pellicle peptides, and the highest scoring control sequence further validate these peptides for enamel interactions (Figure 6). The pattern of half the control sequence scores falling below the range of these peptide scores validates the assumption of non-interaction and supports the hypothesis that regions with pellicle proteins that are not ever observed in the pellicle are evolved to not bind enamel. High scoring peptides are from keratins, calmodulins, cystatins, and others (Supplemental Table 9).

### 4.8 Matthews correlation coefficient plot

The complexity of an MCC curve informs the capacity for improvement: untrained matrices show large local minima, which are lost with improvement (Figure 2c). MCC curves for trained matrices are broader with decreased complexity, suggesting that these are near the end of the respective training paths. The MCC plot also shows the cutoff value with the most discriminative ability.

### 4.9 Comparison to previous work

We extended the methodology for sequence-based prediction of inorganic surface binding peptides to naturally occurring peptides observed in the enamel pellicle. Sampling the amino acid substitution matrix space by selecting among a diverse database proved efficient and useful. As seen previously for artificial phage display derived inorganic surface binding peptides (Oren et al., 2007), amino acid substitution matrix methods can learn contextual patterns, now including physiologic salivary enamel pellicle peptides.

Further understanding of biomineralization proteins and peptides may be gained by considering catalytic activity, structural features, cleavage sites, post-translation modifications, and evolutionary conservation, in the context of the pellitrix scores. While no other tool known to us can learn the patterns in such a small heterogeneous sequence set, the analysis presented here demonstrates the ability of this approach to predict, and therefore interrogate and design protein-hydroxyapatite interactions.

## 5 Conclusions

We demonstrated that enamel pellicle peptides contain subtle sequence similarities that likely encode hydroxyapatite binding mechanisms. With experimental and algorithmic improvements, our substitution matrix-based peptide comparison protocol represented the pellicle peptide similarities in an amino acid substitution matrix (pellitrix) that discriminates pellicle peptides from control sequences with near perfect accuracy (0.99 AUC). We showed that pellitrix can recapture the peptides from their source protein sequences, and that this can be applied as a tool to predict hydroxyapatite interaction regions within relevant proteins. Analysis of relationships between the pellicle peptide sequences indicates that adjacent carboxyl or amide residues facilitate calcium interactions, that alternating hydroxyl moieties mediate phosphate interactions, and that stretches of prolines stabilize extended conformations. This protocol was built as a freely available software suite called Mat4Pep to learn similarities in any set of peptides, for bioengineering design and analysis of any biological function.

## Acknowledgements

We thank Dr. Hector Huang for help with mass spectrometry, and Dr. Bill Landis for help directing revisions.

